# Reappraisal of the Dilution and Amplification Effect Framework: A Case Study in Lyme Disease

**DOI:** 10.1101/2025.01.13.632839

**Authors:** Shirley Chen, S. Eryn McFarlane

## Abstract

The role of biodiversity in regulating zoonotic disease in ecological communities has been broadly referred to as the biodiversity-disease relationship in disease ecology. Whether biodiversity decreases or increases disease risk, known as a dilution or amplification effect respectively, remains unclear. The literature has focused on the strength, generality, nature, and context dependencies that could explain contradictory evidence. We suggest that a continued focus on this approach to resolving the biodiversity-disease debate detracts from a more foundational problem with testing these dilution and amplification hypotheses, in that these hypotheses are not falsifiable as proposed. When tested and interpreted as net effects in a system, these hypotheses do not possess a true null outcome and they are vulnerable to *ad hoc* explanations. To remedy this problem, we propose that biodiversity and disease risk can be modelled as latent variables in multivariate causal models to test specific mechanistic pathways. We present a case study on Lyme disease through a systematic review and meta-analysis, concluding that there is little evidence for a net dilution or amplification effect. We demonstrate how testing these net effect hypotheses falls short of providing robust evidence for its underlying mechanisms. While these hypotheses have previously been helpful in conceptualizing this idea of biodiversity as a potentially protective factor for human health, they need further specificity and clarity to move forwards with greater unity.

## Introduction

### The biodiversity-disease debate

Wildlife populations are responsible for the emergence of over half of the infectious disease events that threaten human health globally (Jones et al., 2008). Links between anthropogenic activity and biodiversity loss have led disease ecologists to converge on one fundamental principle in the field: changing ecological communities affect the transmission of zoonotic (non-human animal originating) diseases (Allen et al., 2017; Estrada-Peña et al., 2014; Glidden et al., 2021; Keesing et al., 2010; White & Razgour, 2020). Following this theoretical basis, two hypotheses have been proposed to suggest that biodiversity loss can either increase or decrease infectious disease risk through what are known as dilution or amplification effects respectively (Johnson et al., 2015; Keesing et al., 2006; Keesing & Ostfeld, 2021; Ostfeld & Keesing, 2000a, 2000b; Schmidt & Ostfeld, 2001).

While these broad biodiversity-disease relationships (hereafter “diversity-disease”) have existed before their applications in zoonotic disease systems (Wolfe, 1985), criticisms have emerged about the simplicity of the dilution and amplification hypotheses in failing to capture the complex mechanisms and dynamic community interactions within these systems.

When proposed as net effects on a system, the ‘dilution effect hypothesis’ suggests that an increase in host diversity limits overall pathogen transmission in ecological communities whereas the ‘amplification effect hypothesis’ suggests that it would increase overall transmission (**Box 1**; Keesing et al., 2006). The contradictory nature of these net effect hypotheses has also prompted debates surrounding the context dependencies, generality, and underlying principles of the dilution effect (Civitello et al., 2015; Halliday et al., 2020; Halsey, 2018; Ostfeld, 2013; Ostfeld & Keesing, 2013; Randolph & Dobson, 2012; Salkeld et al., 2013; Strauss et al., 2015; Wood et al., 2017; Wood & Lafferty, 2013). In attempts to remediate disagreement in the literature, research recommendations abound (Halliday et al., 2020; Johnson et al., 2015; Keesing & Ostfeld, 2021; Kilpatrick et al., 2017a; Rohr et al., 2020; Stewart Merrill & Johnson, 2020), but ultimately affirm the notion that dilution and amplification hypotheses are inherently testable when proposed as net effects.

### The dilution vs amplification debate cannot be resolved as proposed

Conceptual frameworks describing the dilution and amplification effects have been used to visualize how various mechanistic pathways contribute to an overall net change in disease transmission in the community (**Box 1**; Keesing et al., 2006; Kocher et al., 2023; Luis et al., 2018; Wood & Lafferty, 2013).

#### Box 1

Dilution and amplification effects represent two net outcomes of the same framework describing the relationship between biodiversity and disease risk (**Fig. 1**). These net effects result from the cumulation of multiple component effects in a system to primarily mediate the proportion of competent hosts present to support disease transmission. As a result, a change in average community competence drives a net increase or decrease in disease transmission (Keesing et al., 2006; Ostfeld & Keesing, 2000b).

**Figure 1.**
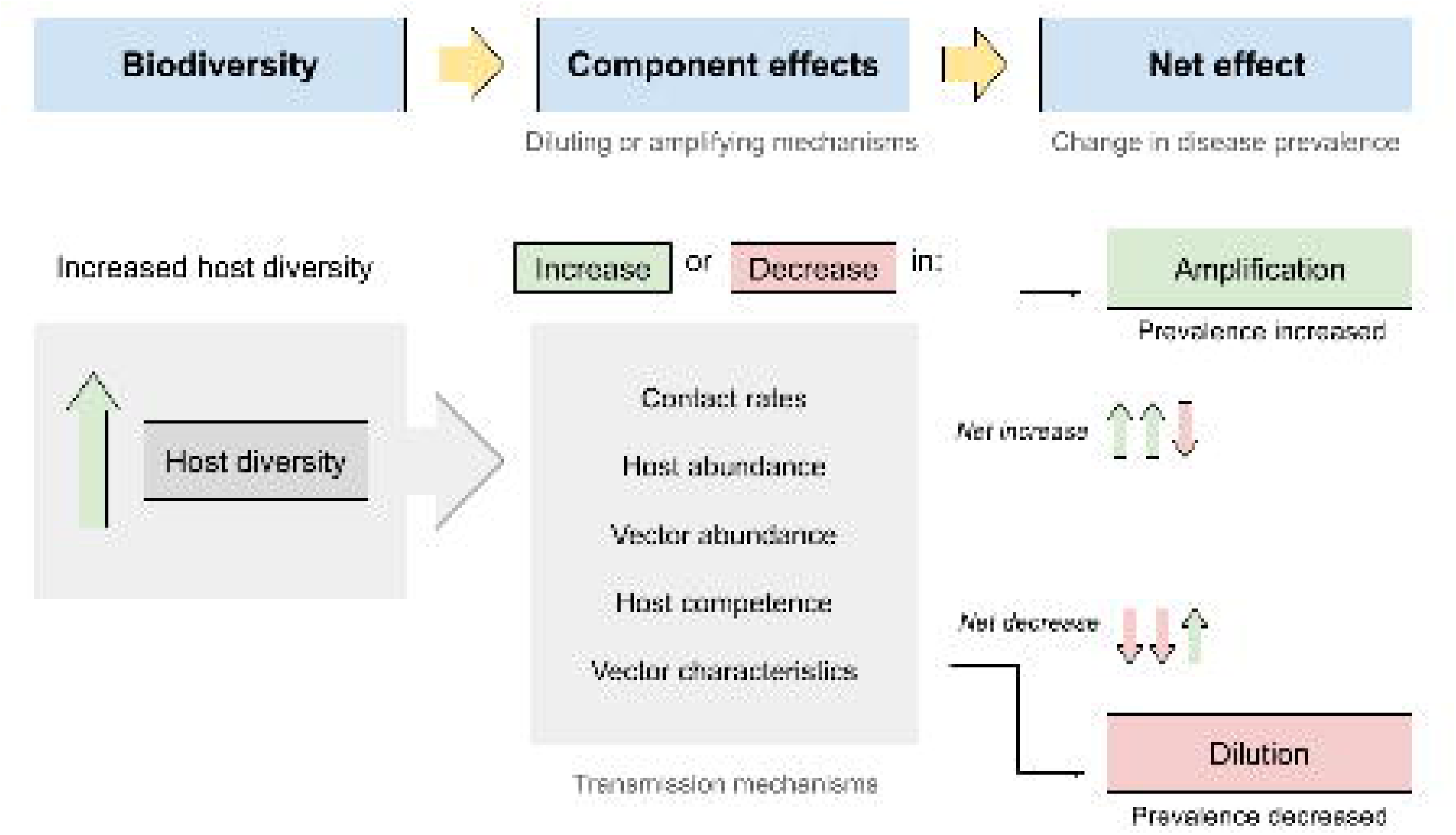
A change in biodiversity, represented specifically by host diversity, can affect the strength of component effects that can increase (amplify) or decrease (dilute) disease transmission. These component effects represent mechanisms operating at a community level and have variable influence on disease transmission. The relative strength of specific mechanisms depends on whether frequency-dependent or density-dependent transmission characterizes the system (Rudolf & Antonovics, 2005). In frequency-dependent transmission, host competence and vector characteristics are more important for promoting transmission, independent of their overall population abundances (i.e. population density). On the other hand, density-dependent systems are more impacted by the population abundances of host and vectors to promote transmission through increased contact rates (Dobson, 2004; Ostfeld & Keesing, 2012). The outcome of all component effects on transmission occurring simultaneously in a system will produce a net increase or decrease in transmission to affect disease prevalence (i.e. disease proportion).

These conceptual frameworks have also been useful to help outline the necessary prerequisites for a dilution effect. Ostfeld & Keesing (2000b) proposed that the following criteria must be met in vector-borne disease systems in order for a dilution effect to be considered: 1) the vector is a generalist such that host diversity correlates positively with feeding opportunities, 2) pathogen transmission is horizontal such that transmission occurs from blood meals only and not between vector generations, 3) host competence varies in the community such that hosts differ in their probability of transmitting the pathogen, and 4) there is a positive correlation between host competence and proportional abundance such that the most competent host is proportionally less abundant when species richness increases in the community. Additional prerequisites have been further supplemented into the literature to specify that diverse communities provide species that reduce the maintenance and transmission capability of amplifying hosts, vectors, or the pathogen itself (Johnson et al., 2015). These prerequisites imply that the driving force for the dilution effect is a change in average community competence with a change in host diversity. Conceptually, this reasoning is logical and general enough that this broader mechanism can be attributed to multiple different, individual causal effects of increased host diversity. However, there is also a tendency towards tautology in this reasoning to suggest that this phenomenon occurs only when the conditions in the system allow it to occur. In these instances, we should be critical about how these hypotheses are being tested and if they continue to be useful in furthering our understanding about diversity-disease relationships.

While conceptual frameworks and interaction webs are useful for generating hypotheses, observing a relationship between host diversity change and *overall* transmission does not lend support for any specific mechanistic pathways occurring. In other words, the dilution and amplification effects, when represented as *net* effects, cannot be tested without first testing their individual components. Others have implied this idea by advocating for a more mechanistic approach to diversity-disease studies (Glidden et al., 2021; Halsey, 2018; Johnson et al., 2015; Keesing & Ostfeld, 2021; Kilpatrick et al., 2017a; Rohr et al., 2020; Stewart Merrill & Johnson, 2020) and some have explicitly tested systems this way (Gandy et al., 2022; Kocher et al., 2023; Strauss et al., 2015). This issue of testing the global effect and backwards inferring potential mechanisms that could be responsible for the observed pattern appears in several prominent diversity-disease studies (Allan et al., 2003, 2009; Brownstein et al., 2005; Ezenwa et al., 2006; Ostfeld & Keesing, 2000a) that have suggested support for dilution or amplification effects. These studies are important to help unveil patterns observed in natural systems, but results should not be extrapolated further to support a singular net effect if component effects have not been explicitly tested. Essentially, patterns are not unequivocal evidence of underlying processes, and should not be interpreted as such (Cooper, 1998). Caution towards this type of causal inference is not novel (Anderson, 2008), and the same principle also applies in the opposite direction of interpretation where evidence for one pathway does not necessarily represent the overall net pattern in the system when other pathways are not observed or accounted for.

Reviews of the dilution and amplification effect framework ultimately have come to contradictory conclusions regarding the strength (Civitello et al., 2015; Salkeld et al., 2013), directionality (Wood et al., 2017), and nature of the relationship (Halliday et al., 2020; Halliday & Rohr, 2019; Johnson et al., 2015; Wood et al., 2017). Context-dependencies exist to potentially dampen the observation of these effects and, additionally, comparisons between studies (Rohr et al., 2020), which leads us to believe that the field should focus its attention on *a priori* approaches to these dependencies and their mechanisms instead of *ad hoc* justifications for contradictory evidence (**Box 2**). The latter may be doing more harm than good when dilution and amplification frameworks are continuously being modified to explain inconsistencies in the emerging data.

### Hypotheses should be unconditionally falsifiable

Controversy surrounding the dilution and amplification effects have historically focused on inconsistencies in how we define disease risk and measure transmission, leading to variable results and overall misconceptions about these hypotheses (Huang et al., 2016; Johnson et al., 2015; Keesing & Ostfeld, 2021; Kilpatrick et al., 2017a; Randolph & Dobson, 2012; Rohr et al., 2020). For instance, disease risk, typically measured as the abundance (i.e. density) or prevalence (i.e. proportion) of infection, has been conflated with parasite diversity (i.e. richness) where the latter does not represent the pathogen transmission *per se* but rather pathogen establishment or colonization (Johnson et al., 2013, 2015). While parasite diversity is not entirely irrelevant from the diversity-disease relationship, it is incorrect to suggest that evidence for host diversity correlating positively with parasite diversity (Dunn et al., 2010; Kamiya et al., 2014) is also evidence in support of the amplification effect. Similarly, if our abundance and prevalence metrics do not equally measure pathogen transmission, which has been suggested to be the case (Huang et al., 2016; Randolph & Dobson, 2012), they are less comparable. These issues are evident and pose valid criticisms regarding the clarity of these hypotheses, however we suggest that a philosophical problem also exists in its foundation: dilution and amplification *net* effects are not falsifiable hypotheses.

### Non-falsifiable hypotheses

At its core, a hypothesis is a scientific model that can offer support for a phenomenon if it can also be rejected through testing (**Box 2**). The net effect hypotheses for dilution and amplification effects cannot be falsifiable if an observed null outcome can also be explained by a cancelling out of opposing component transmission mechanisms. An example for the latter can be observed in the Lyme disease system where deer are both important reproductive hosts for tick vectors while also being low competence hosts for the pathogen. As a result, these two opposing effects, vector amplification and pathogen dilution, have been observed to cancel out and produce a null relationship between deer density and pathogen prevalence (Gandy et al., 2022). While it is possible to test a net effect and definitively observe a null relationship through aggregating known component effects, this requires also testing individual mechanisms as separate hypotheses to be included in a multivariate causal model (e.g. structural equation modelling). That the same pattern (a null outcome) can be explained by multiple processes (either a lack of relationship between diversity and disease risk, i.e. a true null, or cancelling out of amplification and dilution effects by interacting mechanisms) means that the pattern of a null outcome sheds no light on the underlying process.

#### Box 2

##### Falsifiability

A falsifiable hypothesis is a hypothesis that can be empirically tested and which can obtain an observation that disproves the claim (Popper, 2002). Factors that can make a hypothesis non-falsifiable can include issues with specificity (e.g. too vague such that specific predictions are not possible, undefined variables with no clear relationship between them), issues with testing the hypothesis itself (e.g. impossible/supernatural observations, undefined metrics), or issues with the reasoning behind the hypothesis (e.g. circular reasoning, unbounded scope).

##### Ad hoc explanations

An *ad hoc* explanation is when modifications are made to a theory to evade falsification. These modifications may involve introducing *ad hoc* auxiliary hypotheses (i.e. assumptions or prerequisites to a theory), *ad hoc* changes to definitions, and *ad hoc* explanations for inconsistencies in new evidence that would prevent the original hypothesis from being falsified (Popper, 2002). In this case, the hypothesis effectively becomes too flexible to be tested rigorously.

##### Null hypothesis

A null hypothesis is the outcome that describes no effect or relationship between the variables being tested. A good null model considers the “default” system for the relationship being tested (i.e. null predictions should be informed by historical data to create a baseline rather than assuming no relationship existing in the first place). For example, Rohr et al. (2011) critiques the null assumptions about phenological mismatches in testing climate-disease models which assumes perfect synchrony when historical evidence suggests otherwise. In disease ecology studies, specific hypotheses help identify what transmission pathway is being tested, and in turn, what the null model would look like. Whereas the null model for the net effect of biodiversity on disease transmission, more broadly, is less clear due to the ambiguity of this hypothesis and the variations between the disease systems it could be applied to.

Additionally, the broadness of these net effect hypotheses also makes them vulnerable to *ad hoc* explanations of context dependencies (**Box 2**) by using the context as an alternative scenario rather than as evidence for falsification (Clay et al., 2009; Ezenwa et al., 2006; LoGiudice et al., 2008). An *a priori* hypothesis that lacks specificity essentially loses some of its confirmatory nature as a hypothesis as it opens up opportunities for the conflation between hypothesis testing (i.e. does the evidence support my hypothesis?) and *ad hoc* explanations of unexpected results (i.e. how could the evidence still support my hypothesis?; Gelman & Loken, 2013). While this type of *post hoc* reasoning can be arguably justified, since it is often still theory-driven and not spurious, it is also important to note that it also provides explanations of inconsistencies in the data that would otherwise have been evidence to falsify the original hypothesis (Popper, 2002). For example, Ezenwa et al. (2006) observed a negative correlation between non-passerine avian diversity (low competence hosts) and incidence of West Nile virus (WNV) in humans. On the other hand, they observed no correlation between passerine diversity (high competence hosts) and WNV incidence or mosquito infection rates which would have been consistent with the first observation.

Nonetheless, support for a dilution effect was claimed, despite conflicting evidence, by reasoning that these high competence species were more common in all sampling sites. To be clear, *ad hoc* explanations are a systemic issue and we intend to only highlight these subtle lines of reasoning using examples so as to not overlook them as a whole. The concern is that these *ad hoc* explanations may also lend themselves into *a posteriori* hypotheses which at best do not hold strong causal weight (Anderson, 2008). In another example, Clay et al., (2009) observed that rodent species diversity correlated negatively with Sin Nombre virus prevalence in deer mice, suggesting a dilution effect. However, they also note that this outcome could not have been predicted *a priori* because the opposite relationship could have been observed.

The growing list of prerequisites for these dilution and amplification effect frameworks may also unintentionally encourage researchers to look for additional “missing pieces” to unify previously contradicting results regarding the nature of the diversity-disease relationship (Halliday et al., 2020; Wood & Lafferty, 2013). The modification of these prerequisites (“auxiliary hypotheses” *sensu* Popper (2002)) is not inherently problematic as the hypothetico-deductive model of the scientific method follows this process of evaluating tested hypotheses to then modify and generate new testable hypotheses. However, this process also distinguishes between the old and new hypotheses as separate testable hypotheses through a process of modification which should translate into a revision of the system (i.e. ‘strong inference’ (Platt, 1964)). The *ad hoc* modification of auxiliary hypotheses, and indirectly the original hypothesis being tested, should be criticized when it preserves the integrity of the original hypothesis and theoretical framework by changing the conditions of falsification (Popper, 2002). In the case of the dilution and amplification effects, any inconsistencies in the data from these studies should question the support for these net effect hypotheses themselves in addition to their auxiliaries rather than only the latter.

It is tempting to provide *ad hoc* explanations, especially in diversity-disease studies when there is a clear motivation to want to generalize the dilution or amplification effects to different disease systems (Glidden et al., 2021; Halliday & Rohr, 2019; Keesing & Ostfeld, 2021; Ostfeld & Keesing, 2012; Rohr et al., 2020). In the case of the dilution effect specifically, there is a vested interest embedded in finding support for this hypothesis considering how the “biodiversity-buffers-disease” paradigm (Randolph & Dobson, 2012) aligns with the goals of conservation biology. This ‘win-win’ scenario is certainly appealing, but caution should be applied such that this motivation does not lead to the adoption of theories for solely practical reasons. Popper (2002) refers to the idea of conventionalism to highlight the dangers of dogmatism in science, especially when theories remain unfalsifiable as a result of the more insidious practice of *ad hoc* reasoning as opposed to outright denial. To borrow from another philosopher of science, Kuhn (1962) notes that only when the old guard of ideas have been discarded, can there be a scientific paradigm shift.

### Testing mechanistic pathways

Rather than observing an overall pattern and potentially overlooking underlying interactions, identifying evidence in support of specific transmission pathways is crucial for piecing together the bigger picture in the diversity-disease relationship. On the other hand, it is equally important to falsify our initial hypotheses when observing a null relationship instead of generating *ad hoc* explanations as to why amplification or dilution might still be acting.

Clarity and precision in hypothesis generation will allow us to test specific hypotheses in a system, acting as a shield against *ad hoc* explanations. By developing these hypotheses *a priori* (confirmatory) as opposed to continuing to test the whole system and generating *a posteriori* hypotheses (exploratory), we will streamline the progress in disease ecology research and theory development (Anderson, 2008; Grace, 2006). This is especially important to parameterize predictive and causal models which is another goal in disease ecology (Johnson et al., 2015; Rohr et al., 2020) by supplying data about the component effects in the system. As mentioned previously, net effects are testable in this sense if component mechanisms are clearly defined with measurable effect sizes to reflect the proposed transmission drivers in a system. Methods such as structural equation modelling (SEM) to test *a priori* hypotheses in a theoretical framework would be particularly useful in this case where we are essentially examining how a network of direct and indirect transmission drivers influences the overall diversity-disease relationship (Clark et al., 2019; Fearon et al., 2023; Grace, 2006; Grace et al., 2014; Laughlin & Grace, 2019; Shipley, 2000). In this way, we can definitively test and come to conclusions about the existence of any suppressed (i.e. cancelled out) or offsetting (i.e. mediated) effects that are overlooked in net effect hypotheses.

Partitioning our net effect hypotheses into smaller mechanistic hypotheses also addresses practical issues with defining and measuring disease risk. While the hypotheses proposed by Ostfeld & Keesing (2000b) explicitly reference the use of disease prevalence metrics (e.g. proportion of infected vectors or hosts), the use of disease abundance metrics (e.g. density of vectors or hosts) has not been an uncommon representation of disease risk (Huang et al., 2016; Wood & Lafferty, 2013). This problem not only creates difficulties when comparing results across studies but also affects which hypothesis, dilution or amplification, is supported depending on the reference measurement (Gandy et al., 2021; Huang et al., 2016). These contradictory net effect results leave more questions unanswered regarding the underlying interactions between transmission mechanisms and fail to capture the complexity associated with measuring abstract variables with multiple proxy metrics.

As mentioned above, SEM is a useful statistical technique to model casual relationships between intercorrelated variables (Grace, 2006; Laughlin & Grace, 2019; Shipley, 2000). There are two models within the SEM framework that can be applied in this context to conceptualize these relationships: 1) an observed variable model and 2) a latent variable model. We can model component mechanistic relationships as paths between directly measured variables in an observed variable model (**Fig. 2**).

**Figure 2.**
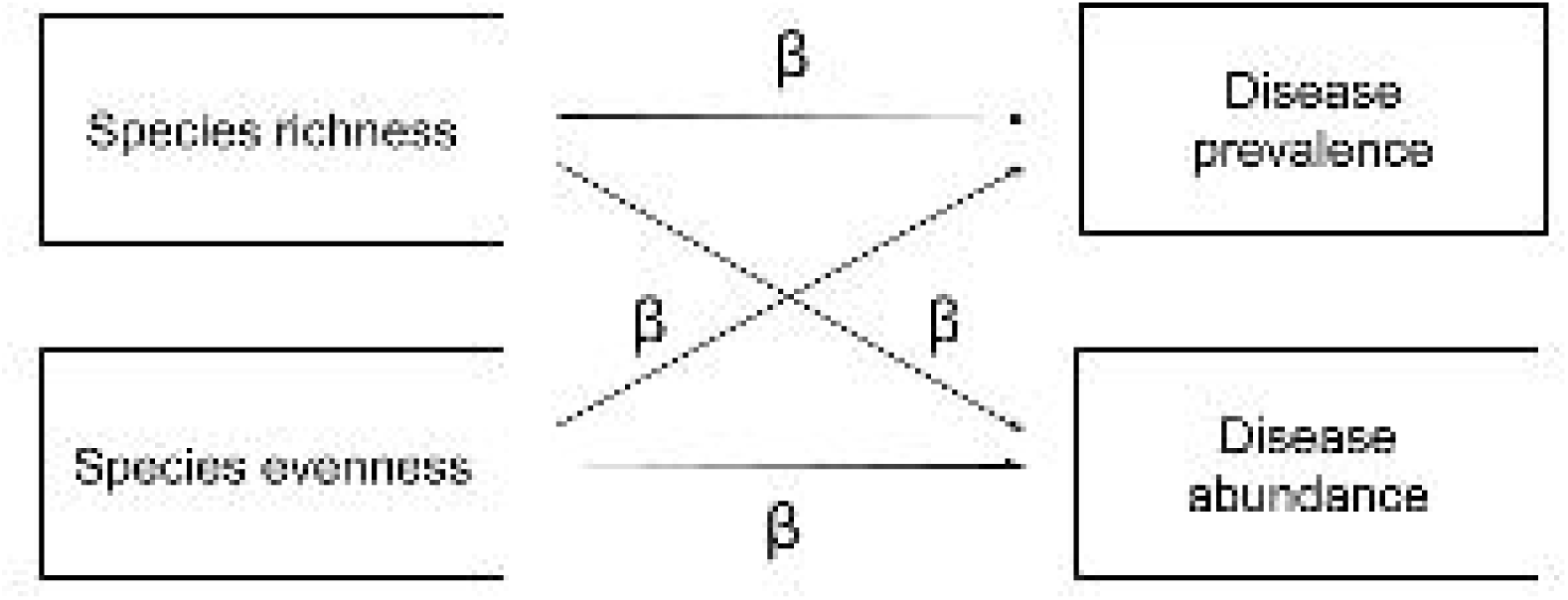
Observed variable model of a simplified diversity-disease relationship that includes only measured variables. Observed variables (species richness, species evenness, disease prevalence, and disease abundance) are represented as rectangles. Single headed arrows represent proposed causal paths between variables with β representing the standardized or unstandardized bivariate regression coefficients.

We can further expand on this observed variable model by thinking of disease risk, and also biodiversity, as unobserved latent variables in a latent variable model (**Fig. 3**). In this way we can visualize the different ways that biodiversity and disease risk can be indirectly measured through proxy metrics in a latent variable model.

**Figure 3.**
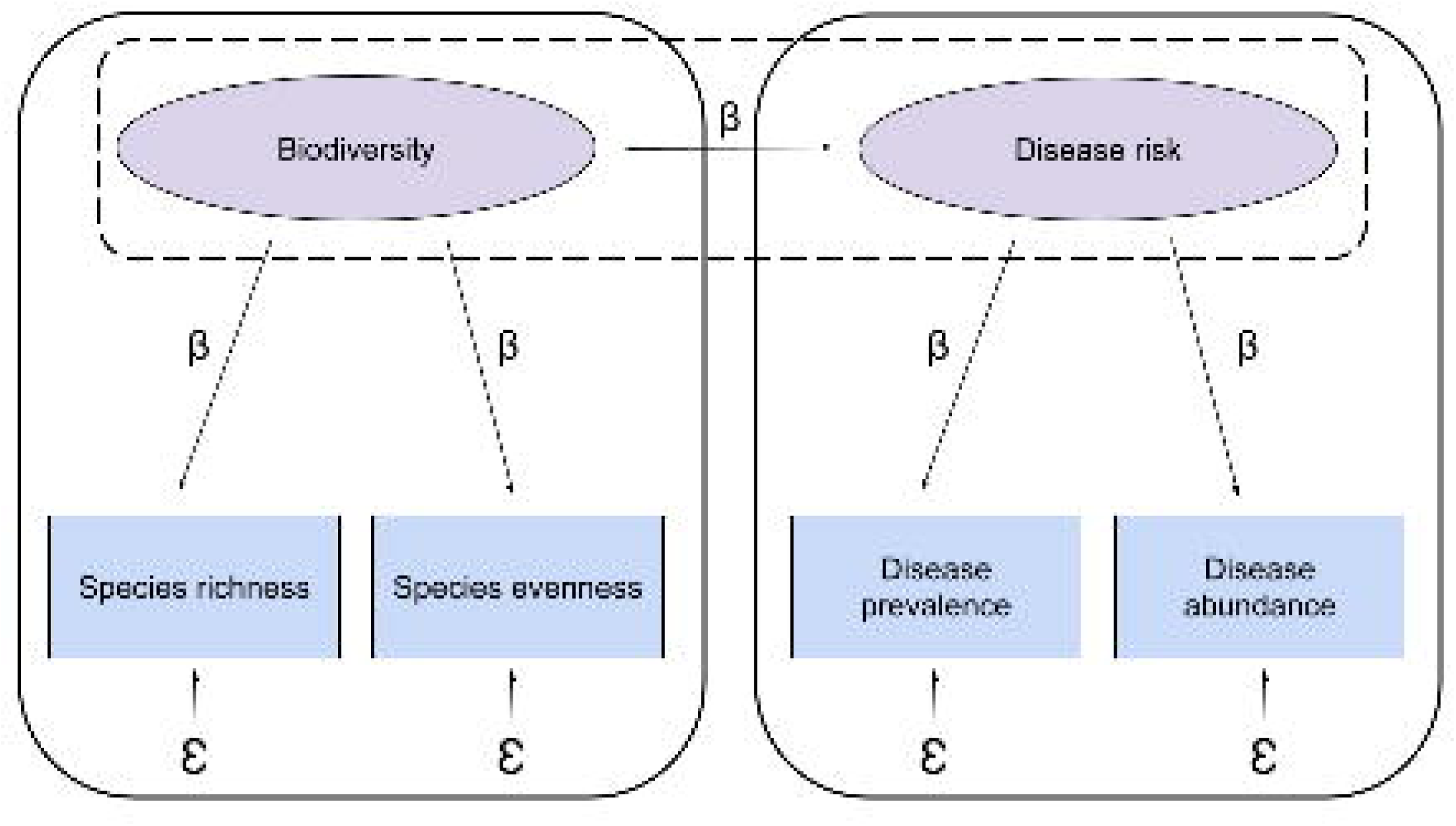
Latent variable model of the diversity-disease relationship with two components: 1) a structural model describing the relationship between latent variables (dashed black outline) and 2) a measurement model for individual latent variables (solid black outline). Latent variables (biodiversity and disease risk) are represented as ovals and observed variables (species richness, species evenness, disease prevalence, and disease abundance) are represented as rectangles. Single headed arrows represent proposed causal paths between variables with β representing the standardized or unstandardized bivariate regression coefficients. [represents error, including measurement error and error that is introduced via stochasticity in the system (i.e. irreducible error).

Latent variable models incorporate a structural model which consists of paths between latent variables. In this case, the path between biodiversity and disease risk represents the hypothesized causal relationship between these two latent variables which are indirectly measured. Mechanistic relationships are also represented by these paths, as was the case in the observed variable model, but the latent variable model makes a distinction between theoretical concepts and proxy metrics being used (Grace, 2006; Shipley, 2000).

When component mechanisms are tested, multiple metrics for biodiversity and disease risk can be tested and assessed for validity in the above model. In other words, we can assess how representative our proxy metrics are of the intended latent concept. Additionally, these multi-indicator models lead to more accurate estimates of the latent construct and greater generalizability of our causal model (Grace, 2006; Shipley, 2000). The advantage of latent variable models is that they consider the reliability of these metrics to estimate measurement error which is inherent to proxy measures. Large amounts of measurement error, indicated by low reliability estimates, can also suggest changes to measurement protocols that may be warranted for future studies (Lamb et al., 2011). As there is still discussion about the best metric to represent biodiversity and disease risk (Huang et al., 2016; Randolph & Dobson, 2012), especially from a practical conservation and human health perspective (Hopkins et al., 2022; Kilpatrick et al., 2017a; Wood et al., 2017), modelling these relationships in a latent variable model can be useful for researchers to gain clearer insight and confidence into the underlying relationships between measured and latent variables within the broader diversity-disease picture.

### A case study: inconclusive net effects in Lyme disease

We present a case study using the vector-borne Lyme disease (LD) system to demonstrate that the amplification and dilution effect hypotheses are not falsifiable. LD is a common and widespread tick-borne disease in temperate regions of North America, Europe, and Asia with a well understood zoonotic cycle and ecology (Kurtenbach et al., 2006). The transmission of the causative bacterial agent, *Borrelia burgdorferi*, is reliant on the proportion of highly competent hosts available to infect ticks, such that a change in their relative abundances would influence the infection prevalence in North American *Ixodes scapularis* ticks (LoGiudice et al., 2003, 2008; Ostfeld & Keesing, 2000a; Schmidt & Ostfeld, 2001). The *Ixodes* tick life cycle is characterized by four developmental stages: egg, larva, nymph, and adult that involve three blood meals where *B. burgdorferi* can be either transmitted from host-to-tick or tick-to-host. Historically, the dilution effect has been the accepted hypothesis in the LD system (LoGiudice et al., 2003; Ostfeld & Keesing, 2000a). However, there has been evidence to suggest that this relationship exists only under specific circumstances which has garnered criticisms about its robustness and generalizability across other LD and zoonotic disease contexts (Kilpatrick et al., 2017b; MacDonald et al., 2022; Randolph & Dobson, 2012; Salkeld et al., 2013; Wood & Lafferty, 2013). Thus, there has been a trend of increasingly mixed support for the dilution effect within the LD literature that continues to be largely unresolved which may indicate that this net effect framework is no longer useful in determining the role of biodiversity in influencing *B. burgdorferi* transmission.

Here, we conduct a meta-analysis and systematic review to evaluate the evidence for amplification and dilution effects in the North American system for LD. However, as the purpose of this study was not to evaluate the evidence for any one specific mechanistic pathway, our hypothesis reflects the dilution and amplification *net* effect framework as proposed in the early literature (Allan et al., 2003; Ostfeld & Keesing, 2000a). That is, if a relationship exists between host diversity and LD prevalence through the regulation of *B. burgdorferi* transmission via multiple component mechanisms, two predictions emerge: a negative relationship would indicate a net dilution effect and a positive relationship would indicate a net amplification effect. It should also be noted that this meta-analysis was conducted naively, prior to a more concrete understanding of the controversies of the dilution effect (Halsey, 2018). As a result, this meta-analysis is moreso intended to highlight issues in the literature that could hamper the ability for future meta-analyses to be conducted and the interpretations of those results.

## Methods

A total of 38 papers were eligible for assessment (out of 526) after conducting a systematic search of the literature with 19 included in the systematic review and 8 in the meta-analysis (**Fig. 4**). A strict selection criteria was implemented to avoid misinterpretations of results due to differential reference metrics. A detailed literature search report, PICO screening framework, and data extraction can be found in the appendix (**Appendix 1 Table 1**).

**Figure 4.**
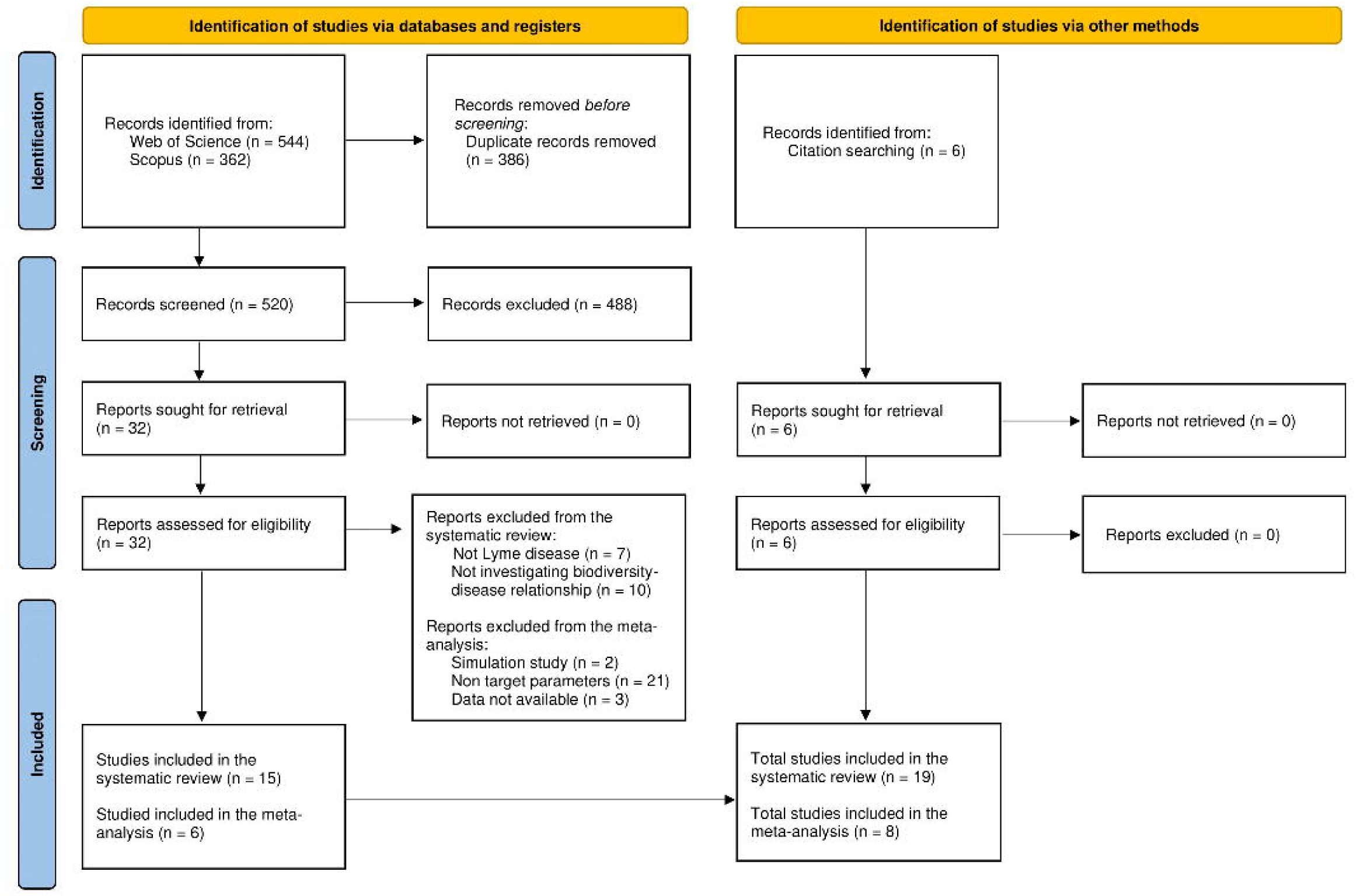
A modified PRISMA flow chart from Page et al. (2021) to include the inclusion process for papers to be included in the meta-analysis. Reports assessed for eligibility (n = 32) from the database searches include those undergone abstract and full-text screening due to a large number of reports excluded (n = 488) during title screening. Studies included in the systematic review (n = 15) and in the meta-analysis (n = 6) are both derived from reports assessed for eligibility (n = 32) from the database searches. Total studies included in the systematic review (n = 19) and in the meta-analysis (n = 8) are the sum of database searches and reports assessed for eligibility from citation searching (n = 6) partitioned into the systematic review (n = 4) and the meta-analysis (n = 2).

**Figure 5.**
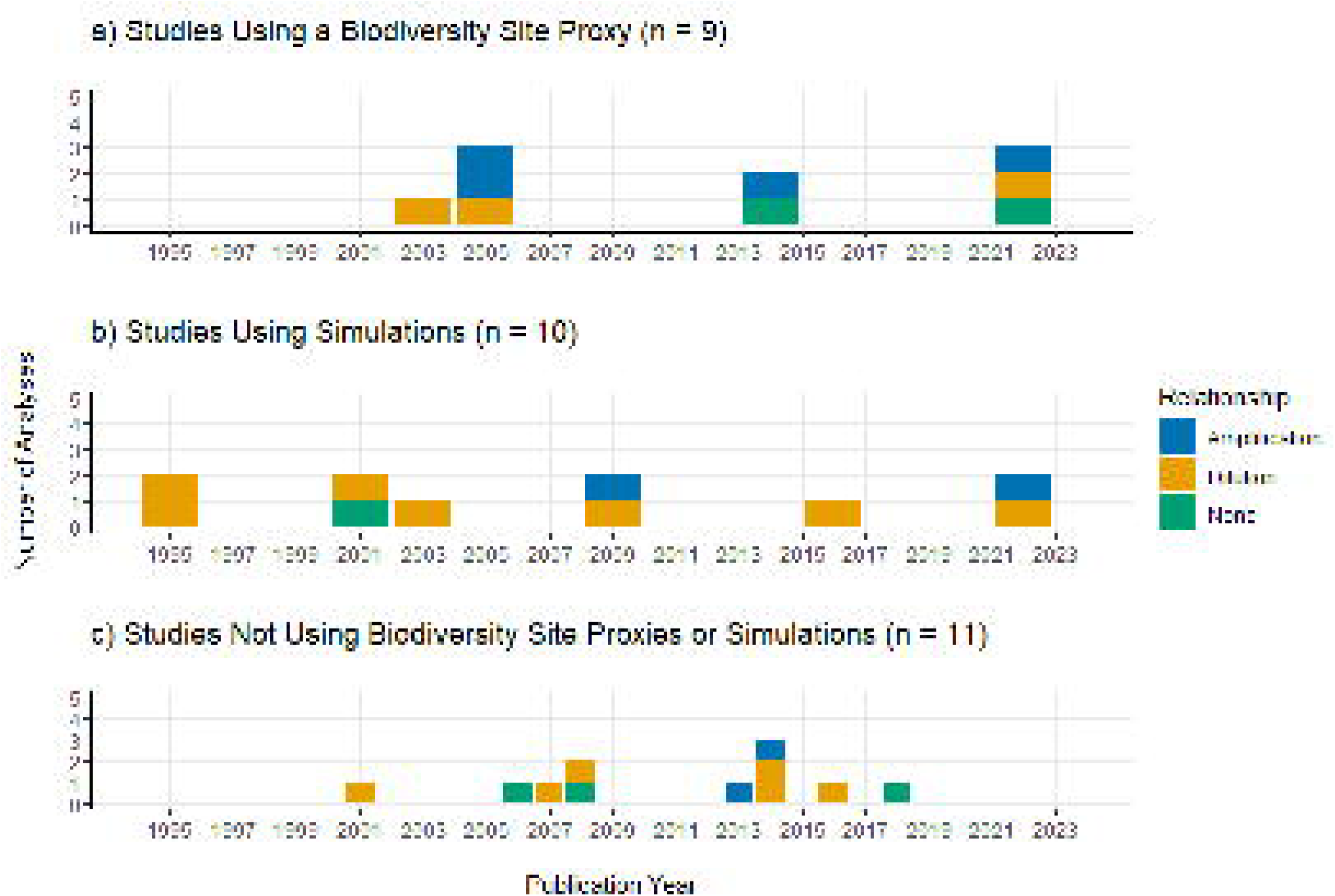
Proportional bar plots showing studies that use (a) site proxies (site size or habitat type), (b) simulations in analyses of the diversity-disease relationship for LD over time. (c) represents studies that did not use site proxies or simulations (i.e. were otherwise empirical studies that used direct biodiversity metrics). The colour coded proportions represent the outcomes (amplification [blue], dilution [orange], none [green]) for each analysis.

### Systematic review

Some papers that we included in the systematic review included multiple separate analyses of the diversity-disease relationship using different parameters. As a result, we categorized each analysis (n = 30) within each paper by the study type (empirical or simulation), biodiversity metric used, LD risk metric used, outcome (dilution, amplification, or none), and primary proposed mechanism. Studies that measured biodiversity using site features such as site size or habitat type (interior vs edge) were categorized as “site proxies” under their biodiversity metric. For detailed information on how each paper was categorized, see the appendix (**Appendix 1 Methods**). We calculated the proportions for biodiversity metric, LD risk metric, and mechanism for each outcome category to show any trends within each effect.

### Meta-analysis

To include test statistics from studies which had the data we were interested in, but which had not tested the specific relationship, we separately conducted statistical analyses for those 3 papers to then include in our meta-analysis (Anderson et al., 2006; Ginsberg et al., 2021; Millien et al., 2023) using logistic regression in R version 4.3.1 (R Core Team, 2023).

Logistic regression was the most commonly used statistical test in the other papers included in the meta-analysis (LoGiudice et al., 2008; States et al., 2014; Werden et al., 2014). We tested small mammal species richness and Shannon Diversity index (H’) separately as independent host diversity variables affecting *B. burgdorferi* infection prevalence. We performed random effect models for all meta-analyses using the {meta} package in R version 4.3.1 (Balduzzi et al., 2019; R Core Team, 2023). A detailed report of the statistical analyses and model building process can also be found in the appendix (**Appendix 1 Methods**).

To examine the effect of the biodiversity metric in the model as a categorical covariate, we performed three separate meta-analyses using: (1) studies using richness or site size (k = 7), (2) studies using H’ or site size (k = 7), (3) studies that did not use site size (k = 6) with richness as the default if both richness and H’ effect size estimates for the same study were available. Studies that use site size as a biodiversity proxy metric for host diversity, under the assumption that increasing site size increases diversity, were identified *a priori* as a significant source of statistical heterogeneity between studies and excluded in this additional meta-analysis 3. Subgroup analyses were not performed due to the lack of available richness, H’, and site size effect estimates for each study and non-independence between some effect sizes (i.e. some studies had more than one effect size derived from the same dataset). For meta-analysis 3, we calculated an additional fixed effects model *post hoc* due to an unexpectedly low I ^2^ heterogeneity reported in the random effects model. This has been attributed to a downwards bias associated with k < 7 studies (Von Hippel, 2015), as a result only the random effects model was included in our results. For fixed model results, see the appendix (**Appendix 1 Results**).

## Results

### Systematic review

Most analyses overall use host richness as their biodiversity metric (n = 16) and NIP (nymphal infection prevalence) as the LD risk metric (n = 9). Within analyses that found an amplification effect (n = 8), richness and site proxies were used equally, and LD abundance was mostly measured. Within analyses that found a dilution effect (n = 16), richness was used the most, and LD prevalence was mostly measured. Within analyses that found no effect (n = 6), combined indices and site proxies were used equally, and LD prevalence was the typical measurement (**Table 1**). Overall, a little under a third of all analyses use site proxies as their biodiversity metric (n = 9) and found mixed evidence for the diversity-disease relationship.

Another remaining third of studies were simulations that did not use a site proxy (n = 10) and mainly found evidence for a dilution effect. Amongst the studies that did not use a site proxy and were not simulations (n = 11), evidence was again mixed (**Fig. 4**).

### Meta-analysis

#### (1) Studies using small mammal richness or site size for biodiversity metrics

Small mammal diversity, measured as small mammal richness or site size, and *B. burgdorferi* infection prevalence were weakly negatively related, indicating a weak net dilution effect, although this was not significant. Across studies (k = 7, n = 108), the pooled effect size in a random effects model is OR = 0.76 (95% CI [0.24, 2.37] using a K-H adjusted t-distribution), t = −0.60, p = 0.571. This means that an estimated 0.76 times increase (i.e. overall decrease) in *B. burgdorferi* prevalence occurs with every unit increase in host diversity, however this cannot be distinguished from a null hypothesis of no relationship (**Fig. 6**). Between study heterogeneity variance was significant (p < 0.0001) and high, estimated to be [^2^ = 1.27 (95% CI [0.24, 7.81]) with I ^2^ = 80.8% (95% CI [61.1%, 90.5%]).

**Figure 6.**
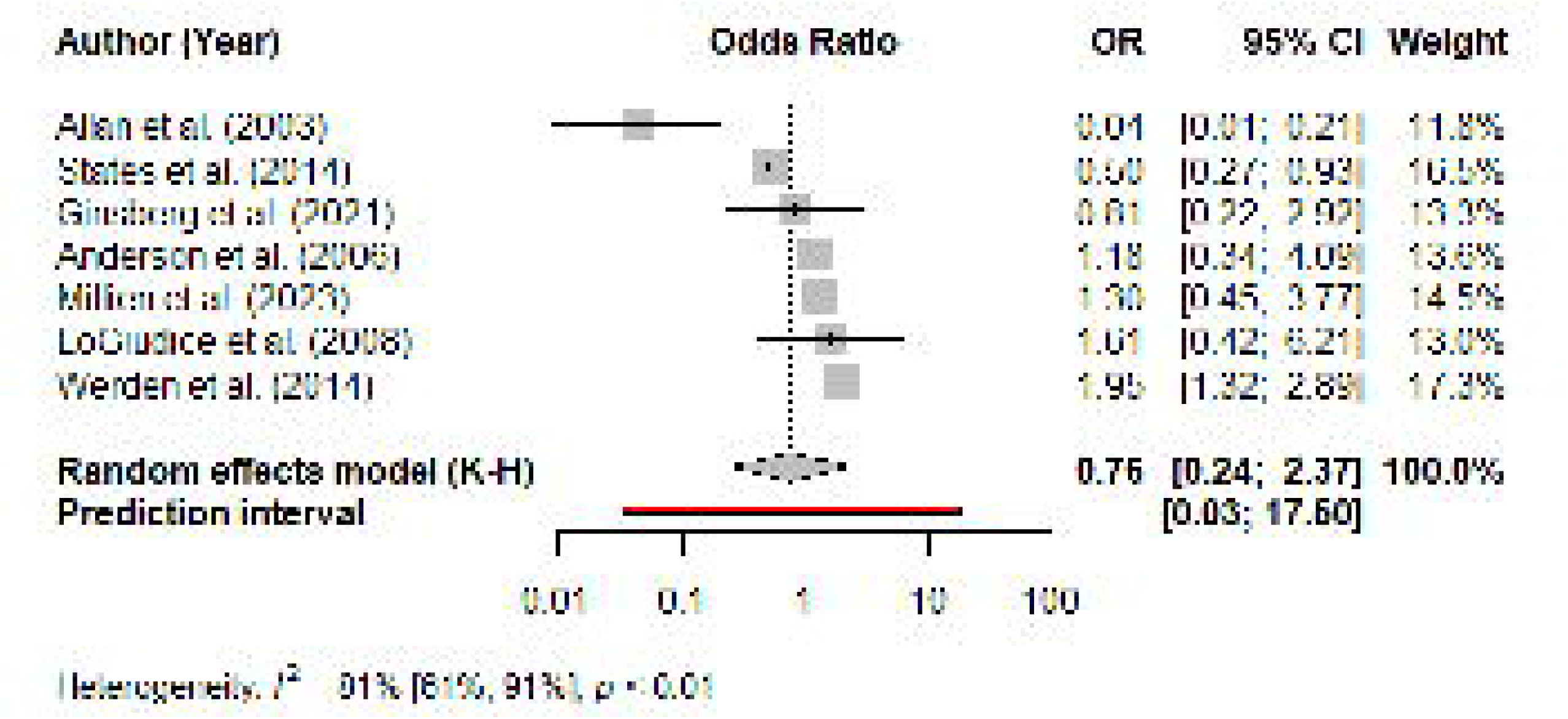
Forest plot showing distributions of odds ratio (with 95% CI bars) for studies describing the relationship between small mammal diversity and *B. burgdorferi* prevalence using richness or site size proxy measures. Grey squares on the CI bars are scaled to size according to the weight of that study in the model. The solid black vertical line represents the null outcome. The diamond represents the pooled effect with the dotted vertical line representing the odds ratio of the random effects model. The red horizontal line represents the prediction interval range given the model results.

#### (2) Studies using small mammal H’ or site size for biodiversity metrics

Small mammal diversity, measured as small mammal H’ or site size, and *B. burgdorferi* infection prevalence were weakly, negatively related, indicating a weak net dilution effect, although this was not significant. Across studies (k = 7, n = 108), the pooled effect size in a random effects model is OR = 0.55 (95% CI [0.13, 2.32] using a K-H adjusted t-distribution), t = −1.02, p = 0.3472. This means that an estimated 0.55 times increase (i.e. overall decrease) in *B. burgdorferi* prevalence occurs with every unit increase in host diversity, however this cannot be distinguished from a null hypothesis of no relationship (**Fig. 7**). Between study heterogeneity variance was also significant (p = 0.0087) and high, estimated to be [^2^ = 1.37 (95% CI [0.10, 9.26]) with I ^2^ = 65.1% (95% CI [21.4%, 84.5%]).

**Figure 7.**
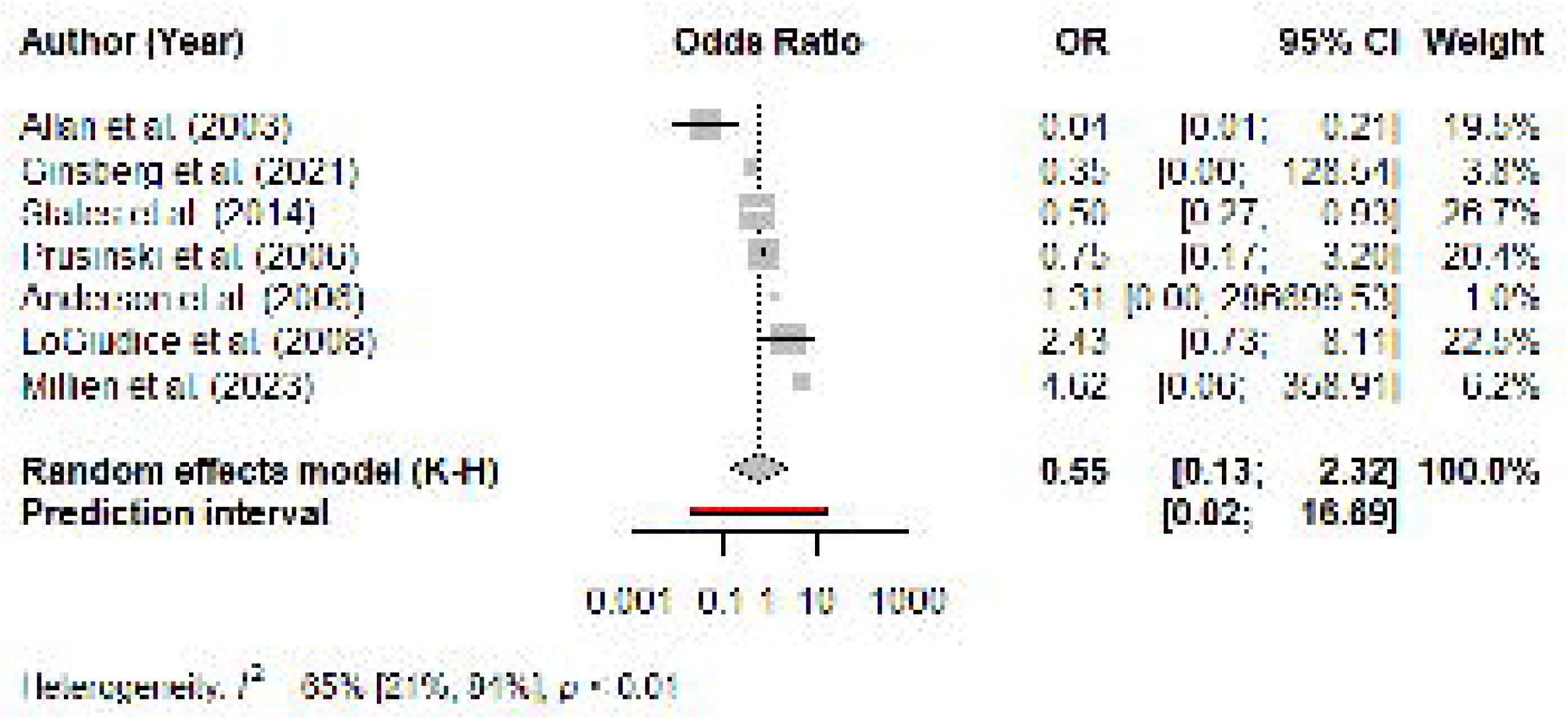
Forest plot showing distributions of odds ratio (with 95% CI bars) for studies describing the relationship between small mammal diversity and *B. burgdorferi* prevalence using H’ or site size proxy measures. The extremely large confidence interval for Anderson et al. (2006) is due to large standard errors from the logistic regression analysis. Grey squares on the CI bars are scaled to size according to the weight of that study in the model. The solid black vertical line represents the null outcome. The diamond represents the pooled effect with the dotted vertical line representing the odds ratio of the random effects model. The red horizontal line represents the prediction interval range given the model results.

#### (3) Studies that did not use site size as a biodiversity metric

Small mammal diversity, measured as small mammal richness or H’, and *B. burgdorferi* infection prevalence were weakly, positively related indicating a weak net amplification effect, although this was not significant. Across studies (k = 6, n = 98), the pooled effect size in a random effects model is OR = 1.48 (95% CI [1.00, 2.20] using a K-H adjusted t-distribution), t = 2.58, p = 0.0494. This means that an estimated 1.48 times increase in *B. burgdorferi* prevalence occurs with every unit increase in host diversity, however this cannot be distinguished from a null hypothesis of no relationship (**Fig. 8**). A test of between study heterogeneity indicated no heterogeneity (p = 0.621), estimated to be [^2^ = 0.05 (95% CI [0.00, 0.56]) with I ^2^ = 0% (95% CI [0.00%, 74.6%]).

**Figure 8.**
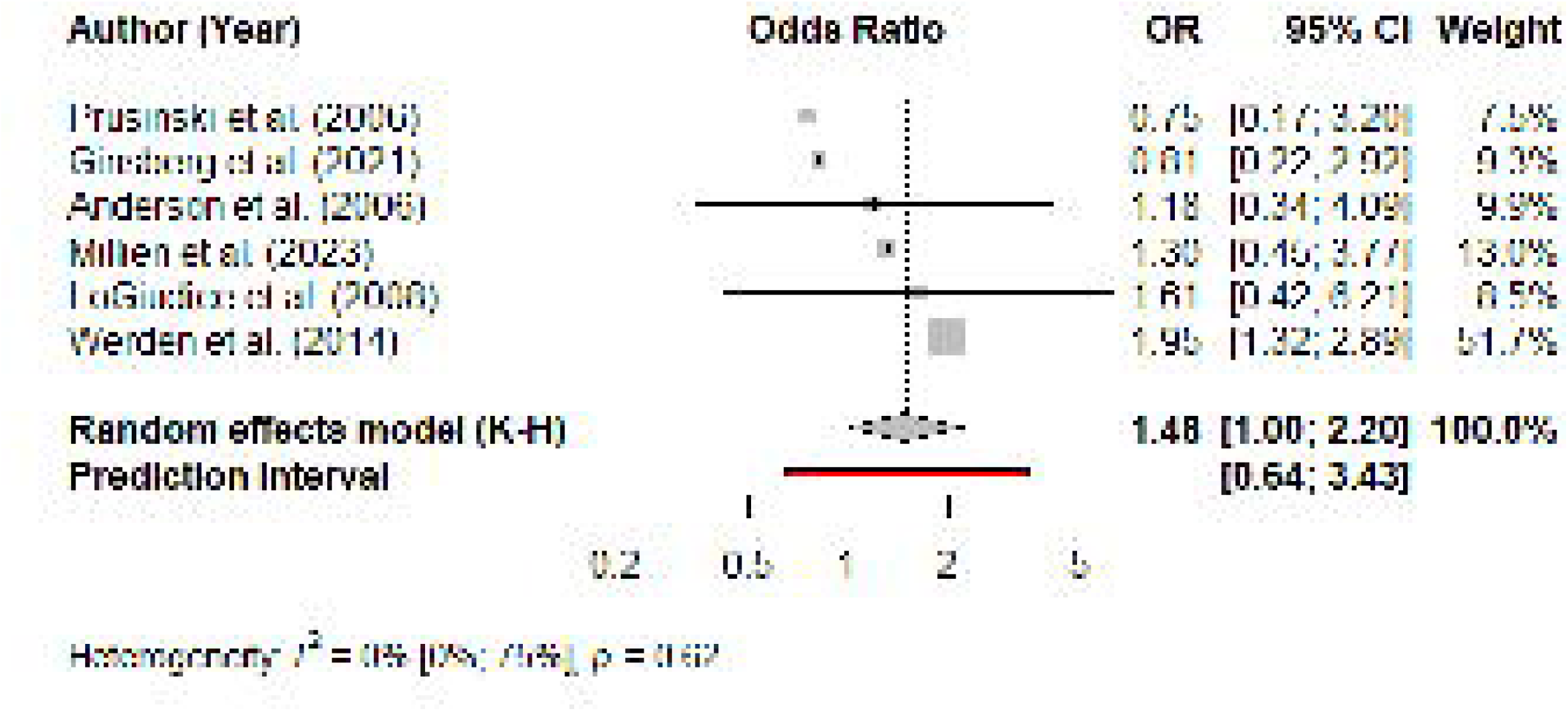
Forest plot showing distributions of odds ratio (with 95% CI bars) for studies describing the relationship between small mammal diversity and disease prevalence using richness or H’ measures. Grey squares on the CI bars are scaled to size according to the weight of that study in the model. The solid black vertical line represents the null outcome. The diamond represents the pooled effect with the dotted vertical line representing the odds ratio of the random effects model. The red horizontal line represents the prediction interval range given the model results.

## Discussion

### Lack of statistical support

Our meta-analyses indicated that there was very weak, non-significant support for net dilution and amplification effect hypotheses for North American LD studies (**Fig. 6, 7, 8**). Most effect sizes were small and non-significant to produce a null global effect, as opposed to being strong in opposite directions, which suggests that there is little reason to believe that future studies measuring this relationship will observe a net effect. No evidence for an observed net dilution or amplification effect can be drawn from our meta-analyses, however this may also be due to minimal information being provided about the specific mechanistic pathways using these particular metrics of biodiversity and disease risk. In other words, strictly quantitative measures of host diversity may not be adequate to represent mechanisms, such as host encounter abundance, that regulate the transmission of *B. burgdorferi* (Linske et al., 2018). A previous meta-analysis investigating this net effect across multiple disease systems, including LD, suggested a similar conclusion where the diversity-disease relationship is inconsistent, with weak effect sizes across studies (Salkeld et al. 2013). This is a null outcome suggesting that cumulative effect of biodiversity on disease risk is specific to the ecological processes and dynamics acting within different disease systems. As a result, our meta-analysis and that of Salkeld et al. (2013) are inconclusive rather than confirmatory of any underlying diversity-disease relationships. In other words, if the individual studies included in a meta-analysis are not explicitly testing the same mechanism, meta-analyses on these studies are not truly as informative as have been interpreted. A different meta-analysis concluded a strong net dilution effect across multiple disease systems but acknowledged the inability of their meta-analysis to test its mechanisms (Civitello et al. 2015). The nature of our initial hypotheses for the present meta-analysis reflects the testing of net effects in the dilution and amplification framework (Keesing et al., 2006). The presence or absence of a net effect is testable in this way; however its component mechanisms are not. This is also meant to highlight the difficulties associated with testing a net effect in a simplistic bivariate model. It lacks the explanatory power of multivariate models, and we suggest that if one is to be conducted, future meta-analyses should focus their attention towards testing the pooled effect of specific mechanistic relationships (Heckley & Becker, 2023; Murray et al., 2019; Vicente[Santos et al., 2023; Young et al., 2013). For example, such a mechanism could be the effect of the relative abundance of *Peromyscus spp.* (deer mice) on nymphal infection prevalence with *B. burgdorferi* (Bouchard et al., 2013; LoGiudice et al., 2008; Mason et al., 2022; Werden et al., 2014). This example would provide information about the strength of the amplifying potential of *Peromyscus spp.* in the LD system across studies, and hence a specific mechanistic relationship that can be evaluated.

The results also indicated that when excluding studies using site size as a proxy metric for host diversity, the direction of the relationship could change (**Fig. 8**). This suggests that some feature of the broader site may drive a decrease in *B. burgdorferi* prevalence, however this should not be attributed to the effects of smaller scale host diversity on disease transmission. Doing so would result in an ecological fallacy – when data at a larger scale is used to infer features of its components (Salkeld et al., 2013). As expected, the heterogeneity decreased very significantly when site proxy was controlled for (I ^2^ = 0%). However, this decrease could be less significant when accounting for an average 28% downwards bias of I ^2^ estimates when k < 7 and true heterogeneity is high (Von Hippel, 2015). We recognize that our meta-analyses suffer from low statistical power due to a small number of studies creating a bias for non-significant results (Deeks et al. 2023). While adjustments were made during model construction to account for this, we nevertheless proceed cautiously with our interpretation. However, we believe it is still useful to note the usage of site proxies beyond the scope of our meta-analysis and view them with a more critical lens moving forwards.

### Latent variable problem

The usage of different metrics of biodiversity and LD risk appear to change the direction of the net effect, despite a lack of statistical support for either effect (**Fig. 6, 7, 8**). While it is tempting to assume that the usage of different biodiversity metrics (e.g. site proxies) is the mechanism for this directional change, we suggest that inconsistent metric usage acts as a red herring for the simpler conclusion. That is, regardless of the metric being used, there is no net effect. As discussed previously, differential conclusions regarding the diversity-disease relationship have been due in part to the usage of different reference metrics. Huang et al. (2016) highlights this problem in studies on vector-borne disease systems more broadly to criticize the ambiguity of the dilution effect. In this case, the driving mechanism for the dilution effect does not account for an increase in total vector abundance in these systems, due to an overall increase in feeding opportunities, that occurs concurrently with a decrease in infection prevalence with feeding diversion. These two ‘abundance’ and ‘prevalence’ metrics to measure disease risk are both used commonly in LD research (Wood & Lafferty, 2013). In our review of the literature, most analyses that observed an amplification effect measured disease risk as LD abundance whereas most analyses that observed a dilution effect used LD prevalence (**Table 1**). While this pattern exists, it should also be noted that this inconsistent metric usage makes it difficult to dissect out the true underlying diversity-disease relationship from noise created due to differences in metrics. Should it be the case where we standardize our metrics, evaluating the relationship will become more clear. However, as discussed previously, there is disagreement on the most representative metric (Hopkins et al., 2022; Huang et al., 2016; Kilpatrick et al., 2017a; Randolph & Dobson, 2012; Wood et al., 2017). In essence, there is a problem with poor or incomplete representation of the latent variables, biodiversity and disease risk, in diversity-disease studies, obscuring our ability to make comparisons across studies that come to contradictory conclusions. This “apples-to-oranges” comparison problem is made clear during the establishment of an inclusion criteria for studies in a meta-analysis which results in a trade-off between statistical power and validity in the final interpretation. For example, a broader inclusion criteria that includes studies measuring biodiversity and disease risk in multiple different ways would allow for more studies to be included, and thus having a greater sample size. However, this would also constrain our ability to interpret the results as evidence for any one specific mechanism driving the overall effect since multiple metrics represent multiple different mechanisms that are pooled together. Additionally, while we cannot know if single metrics being reported are due to preferential reporting, studies that evaluate and report multiple metrics of these latent variables can help visualize instances of multiple pathways concurrently increasing and decreasing disease risk in the same system (Gandy et al., 2021, 2022; Kocher et al., 2023). Insight into how these component effects may cancel out a net effect allows us to make more informed interpretations regarding a null effect when more information is provided in these multivariate analyses.

### Context dependencies

One major criticism regarding the generalization of the dilution effect in LD focuses on the assumptions made about the system that would suggest the effect to be present. In a way, the dilution effect hypothesis uses overall host diversity as a proxy for the proportion of highly competent hosts when assuming a negative correlation (Ostfeld & Keesing, 2000b). For example, *Peromyscus leucopous* (white-footed) mice are commonly regarded as the primary reservoir host of *B. burgdorferi* in Eastern North America due to their high competence, abundance, and conspicuousness. In depauperate communities, *P. leucopus* are expected to persist which contributes to their amplification potential as hosts in low species diversity settings (Ostfeld & Keesing, 2000a). However, inconspicuous, and scarcely studied, reservoir hosts populations of *Blarina brevicauda* (Northern short-tailed) and *Sorex cinereus* (masked) shrews have been identified as being equally if not more important predictors of LD risk (Brisson et al. 2008). This claim threatens one of the existing assumptions of the dilution effect hypothesis, that the most competent host should be the most abundant in depauperate communities (Ostfeld & Keesing, 2000b), and introduces sampling biases when testing the framework. Similarly, the use of site proxies assumes that with increasing site size and at habitat edges, host diversity increases (Allan et al., 2003; Brownstein et al., 2005). These assumptions do not always hold true as the lack of a relationship has been observed (Linske et al., 2018; Mason et al., 2022), which means the true relationship between host diversity and LD risk could differ from what has been reported in the literature. Furthermore, another central assumption that *I. scapularis* is a true generalist has also been observed to not be the case (Ginsberg et al., 2021; Goethert & Telford, 2022), which threatens the assumption about the vector’s generalized host feeding patterns (Ostfeld & Keesing, 2000b). As a result, transmission dynamics may be mediated by the behaviour of ticks themselves rather than the available proportions of high competence hosts. This is an example of where these assumptions function as auxiliary hypotheses to the dilution effect framework in the LD system such that evidence that falsifies these auxiliaries should also weaken support for the primary framework. However, when interpreting unexpected results while testing the dilution or amplification effects, caution should be exercised as to not make *ad hoc* suggestions that question the validity of auxiliary hypothesis as a substitute for falsifying the framework. For example, Werden et al. (2014) observed that the assumption that highly competent mice would dominate depauperate communities did not hold up in all of their sampled sites.

However, they suggested that this represents the context-dependent nature of the biodiversity-disease relationship. In this way, one of the auxiliaries can be falsified but not the dilution effect itself. When we observe a null outcome of a net effect, it may not be clear which hypothesis lacks support in these frameworks that include multiple auxiliaries. As a result, it is increasingly important for auxiliaries to be tested independently and rigorously before including them in these dilution and amplification frameworks to allow for accurate interpretations of observations and falsification of these frameworks directly.

Additionally, a significant proportion of the diversity-disease LD literature is reliant on simulation studies that are modelled off these assumptions about the system (**Fig. 5**). Two of such assumptions necessary to produce a dilution effect in these models are 1) the decrease in proportion of highly competent hosts when host diversity increases, and 2) an increase in tick feedings on low competence hosts due to increased contact rates (Ogden & Tsao 2009). Both assumptions have been falsified with empirical evidence (Ginsberg et al., 2021; Goethert & Telford, 2022; Linske et al., 2018; Mason et al., 2022) which reduces the validity of these simulations in generalizing the dilution effect across all LD systems. If the dilution effect only holds in extremely specific circumstances, then it should not be considered as a foregone conclusion that it will be observed. A null outcome must be considered in empirical studies. Simulations can be used to demonstrate the possibility of processes underlying patterns in empirical systems, but they cannot be used to justify the possible existence of patterns that have not been documented. Less specific models with relaxed assumptions produce both dilution and amplification effects which further demonstrates the inconclusiveness of testing for net effects in this system (Occhibove et al., 2022; Ogden & Tsao, 2009).

### How can future studies improve research on the biodiversity-disease relationship?

A mechanistic focus on the diversity-disease relationship entails a bottom-up approach to investigating component mechanisms individually as part of a larger network. Working to develop and test specific, *a priori* hypotheses that describe a clearly defined mechanism linking biodiversity to disease transmission allows us to evaluate the effect of individual pathways and their interactions with other pathways. Additionally, it allows us to more readily accept that not all proposed mechanisms will be statistically or casually related to disease risk, which confers an advantage over testing a net effect framework. One argument for the practical usage of net effects is that they provide information about the cumulative increase or decrease of disease risk in a community, and as a result we can gauge the net impact of biodiversity conservation interventions (Kilpatrick et al., 2017a). However, these net effects being a cumulation of multiple different pathways means that any given interventions can have potential trade-offs. When these context dependencies exist in a system, there can be positive and negative effects on disease risk occurring simultaneously as a result of a change in biodiversity. Increasingly targeted strategies can minimize off-target effects; however, these require robust evidence at the mechanism level rather than that of a net outcome. This evidence gap in our mechanistic understanding and the inconsistency of these relationships under different circumstances hinders our ability to translate diversity-disease research into practice when the risks outweigh the potential benefits (Hopkins et al., 2022).

In this way, we advocate for a paradigm shift moving forwards by placing less emphasis on net effect generalizations and by no longer framing the diversity-disease relationship as a static, one effect size phenomena occurring in a system. These net effect approaches to the dilution and amplification effect framework are less helpful than they might have originally been in the past when exploring this emerging relationship between diversity change and disease transmission in the literature. Since their proposal, these preliminary hypotheses have generated new, mechanistic hypotheses that we ought to lead with in future studies (Allan et al., 2010; Burkett-Cadena et al., 2021; Gandy et al., 2021, 2022; Ginsberg et al., 2021; Kocher et al., 2023; Levi et al., 2012; Levine et al., 2016; Linske et al., 2018; MacDonald et al., 2022; Young et al., 2017). Furthermore, by accepting that these hypotheses can be falsified, using more specific tests (such as SEM) to assess component effects and evaluate the potential for latent variables, we believe that we can learn more about the potential for a relationship between biodiversity and disease.

---

**Table 1.** Proportion of individual analyses (n = 30) from studies (n = 19) in the literature suggesting an amplification, dilution, or no effect of biodiversity on LD risk. Within each outcome, the proportion of different biodiversity metrics and LD risk metrics measured are shown. Combined indices metric includes species diversity indices that evaluate evenness and richness such as the Shannon Diversity index. Site proxy metrics include site size or habitat type (interior vs edge). LD abundance metric represents DIN (density of infected nymphs) only. LD prevalence metric represents NIP (nymphal infection prevalence), adult ticks, or hosts. Tick burden represents the density or total number of feeding or questing ticks.

| Metric |  | Amplification | Dilution | None |
| --- | --- | --- | --- | --- |
| Biodiversity | Species richness | 50.0% | 68.8% | 16.7% |
|  | Species evenness | 0% | 0% | 16.7% |
|  | Combined indices | 0% | 12.5% | 33.3% |
|  | Site proxy | 50.0% | 18.8% | 33.3% |
| LD risk | LD abundance | 37.5% | 18.8% | 33.3% |
|  | LD prevalence | 25.0% | 50% | 66.6% |
|  | Tick burden | 25.0% | 18.8% | 0% |
|  | Human incidence | 12.5% | 12.5% | 12.5% |

## Data Accessibility Statement

All data and code for this project can be found at https://github.com/scjpg/lyme-biodiversity-biol4000.

## Competing Interests Statement

The authors declare no conflicts of interest.

## Author Contributions

Shirley Chen and S. Eryn McFarlane conceptualized the project. SC determined the methodology for the systematic review and meta-analysis, under the guidance from SEM. SC did the systematic review and all analyses for the meta-analysis. With input, editing and feedback from SEM, SC wrote the initial draft of the manuscript. Both authors have edited this manuscript.

## Acknowledgements

The authors are grateful to Robert Tsushima, Alex Mills, Chris Lortie, William Jones, Murielle Ålund, Mario Zuliani, Jenna LeBlanc, Liz Mandeville and Beth Clare for feedback on this manuscript. This manuscript is the result of SC’s undergraduate thesis project at York University.

## A Appendix 1

### METHODS

#### Database search

We conducted a systematic search following the Preferred Reporting Items for Systematic Reviews and Meta-Analyses (PRISMA) guidelines (Moher et al., 2009). Using the Web of Science and SCOPUS databases (1900–Present), we conducted the original search string: “mammal* AND tick* AND disease* AND (dilut* OR amplif* OR biodiversity OR diversity OR richness OR evenness) NOT (livestock OR domestic*)” (November 2023). We obtained a total of 447 papers after removing duplicates. We conducted another search due to a low output of final papers in the original search string with an updated search string: “mammal* AND lyme AND (dilut* OR amplif* OR biodiversity OR diversity OR richness OR evenness) NOT (livestock OR domestic*)” (December 2023) to obtain an additional 73 papers after removing duplicates. We conducted additional citation searches to further obtain 6 papers for a total of 526 unique papers overall.

For the systematic review, we included papers if they tested the biodiversity-disease relationship with respect to LD explicitly or in *Ixodes* tick hosts of *B. burgdorferi* (n = 19). We included both empirical and simulation studies using multiple different biodiversity metrics and disease metrics. To screen papers for the meta-analysis, we established a criteria following a modified version of the PICO framework to establish a target population, intervention, and outcome for our research question (**Table 1**) (Miller & Forrest, 2001).

**Table 1.** PICO framework to develop a screening criteria for the meta-analysis.

|  |  |
| --- | --- |
| **Population** | Small mammal hosts and *I. scapularis* ticks |
| **Intervention** | Change in small mammal host diversity (richness, $H'$, or site) |
| **Comparison** | Not applicable in this case |
| **Outcome** | *B. burgdorferi* infection prevalence in ticks or small mammal hosts |

We restricted the target study site location to Eastern North America north of the 37° N latitude due to behavioural differences in southern *I. scapularis* ticks (Ginsberg et al., 2021).

We included papers that met the criteria and reported an effect size and standard error (n = 2) or sufficient test statistics to calculate both (n = 3). Additionally, we included papers that did not directly test this relationship but provided data to run the appropriate statistical analyses independently (n = 3). We contacted the authors from 5 papers to request raw data for independent analysis and received data back from 2. A total of 8 papers were included in the final selection.

#### Data extraction

For the systematic review, we assessed studies based on the biodiversity and disease metric used, proposed mechanisms, relationship outcome, study type, and publication year.

Individual analyses (n = 30) from all studies (n = 19) were categorized by the following extracted information: study type, biodiversity metric, disease metric, relationship, and mechanism.

For the meta-analysis, we extracted sample size, effect size, standard error, and total sampling area (ha) from each paper (Allan et al., 2003; LoGiudice et al., 2008; Prusinski et al., 2006; States et al., 2014; Werden et al., 2014). Effect sizes and associated standard errors were extracted directly or calculated from extracted test statistics and transformed into log odds ratio (“logit”) using the {esc} package in R version 4.3.1 (Lüdecke, 2018; R Core Team, 2023). We also extracted raw data from papers that did not test the specific relationship between small mammal diversity and *B. burgdorferi* prevalence (n = 3) to conduct our own independent analyses and obtain an effect size and standard error.

### Raw data extraction

#### Millien et al. (2023)

All sampled sites were included for n = 29 total samples. We extracted small mammal species and trap count data to manually calculate richness and H’ using the {vegan} package in R version 4.3.1 (Oksanen et al., 2022; R Core Team, 2023). We extracted data for the total number of questing nymphs and number of infected nymphs to calculate NIP. We contacted the author to provide site data and manually calculated the total sampling area.

#### Ginsberg et al. (2021)

Only the WI, MA, and NJ sites were included due to the latitudinal boundary constraint for n = 3 total samples. The RI site was within the appropriate boundary but excluded from our analysis because the authors indicated that full small mammal sampling did not occur at this site. We extracted species and trap count data for small mammals only as larger mammals and birds were included in the raw dataset. Small mammal designations were identified in the paper and cross-validated with small mammal species from other papers included in the meta-analysis. Data was extracted from the “original codes” table or “corrected codes” when applicable. Richness was manually calculated and H’ was calculated using the {vegan} package in R version 4.3.1 (Oksanen et al., 2022; R Core Team, 2023). *Sorex spp.* was counted as one species in our analysis. We extracted data for the total number of questing adults and number infected adults to calculate AIP because nymphs or larva were not collected. Only a subset of the total adult ticks collected were tested for infection. Total sampling area was manually calculated using the dimensions and numbers of arrays at each site.

#### Anderson et al. (2006)

Only county-level data was provided for the 96 sites sampled so only counties were included (Anne Arundel, Calvert, Charles, Prince George’s, St. Mary’s) for n = 5 total samples. We contacted the author to provide small mammal species and trap count data to extract and we manually calculated richness and H’ was calculated using the {vegan} package in R version 4.3.1 (Oksanen et al., 2022; R Core Team, 2023). We extracted data for the total number of small mammals and number of infected small mammals to calculate HIP. Only a subset of the total small mammals trapped were tested for infection (*Peromyscus leucopus*, *Mus musculus*, *Microtus pennsylvanicus*, *Blarina brevicauda*). Total sampling area was manually calculated using the site area provided.

**Table 2.**
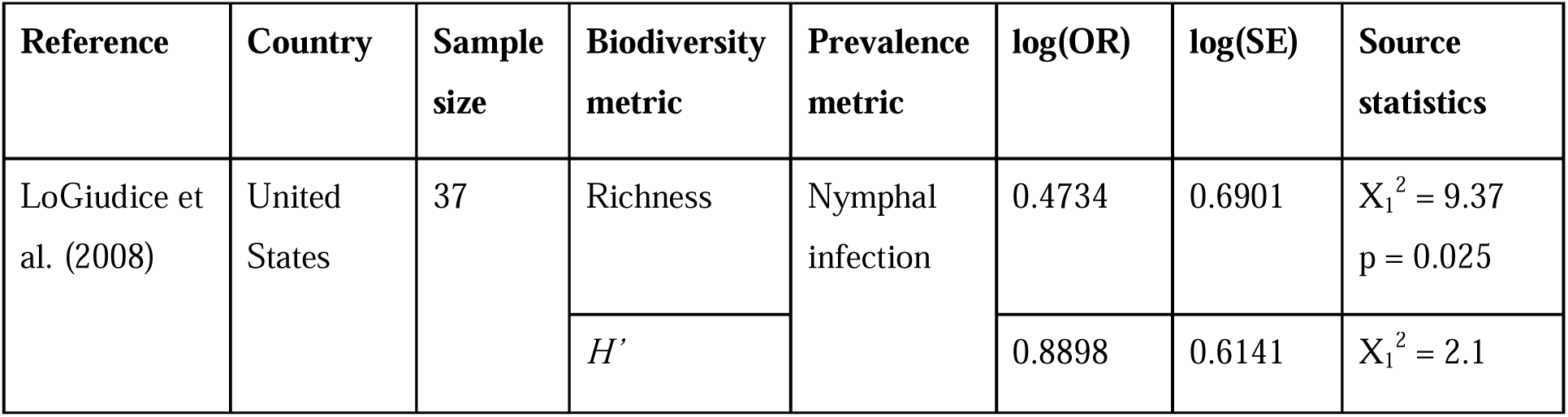
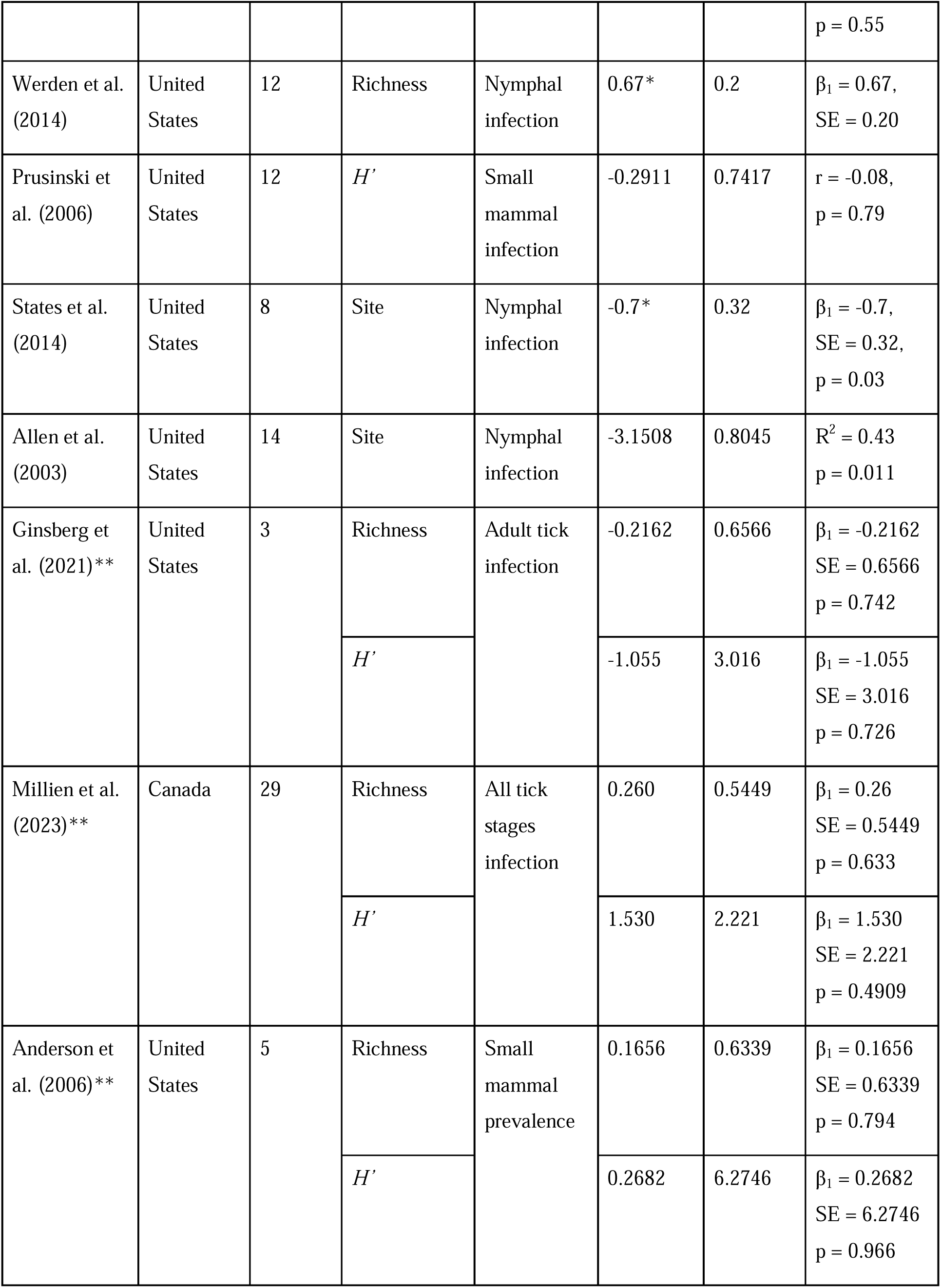
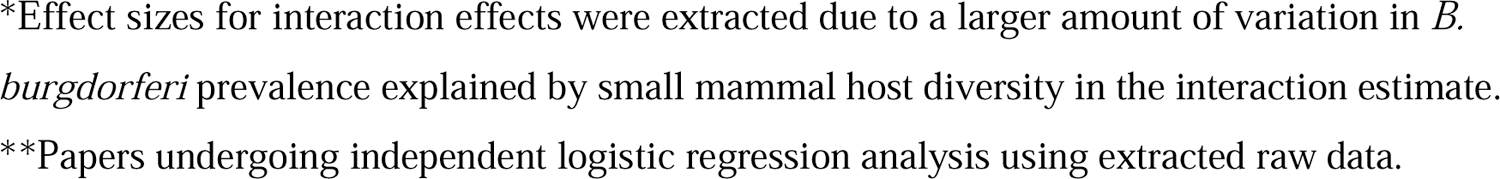
Studies included in the meta-analysis and extracted information: country, sample size, biodiversity metric, disease prevalence metric, effect size, standard error, and source statistics from the original analysis.

### Systematic review

Some papers included in the systematic review included multiple separate analyses of the biodiversity-disease relationship using different parameters. As a result, we categorized each analysis (n = 30) within each paper by the study type (empirical or simulation), biodiversity metric used, LD risk metric used, outcome (dilution, amplification, or none), and primary proposed mechanism. We calculated the proportions of biodiversity metric, LD risk metric, and mechanism for each outcome category to show any trends within each effect.

### Meta-analysis

A Sidik-Jonkman estimator was used to estimate the heterogeneity variance, □^2^, which accounts for high expected heterogeneity and a small number of studies in the model (Sidik & Jonkman, 2007). Additionally a Knapp-Hartung (K-H) adjustment was applied to further reduce Type I error associated with limited studies with binary outcomes and varying sample sizes by broadening confidence intervals (IntHout et al., 2014). We measured heterogeneity using both □ ^2^ and I ^2^ to reduce bias associated with statistical power in limited study meta analyses that is associated with the traditional Cochran’s Q statistic (Harrer et al., 2021; Higgins & Thompson, 2002).

We used a contoured funnel plot to visualize the presence of publication bias due to small-study effects with studies using richness as the default metric (k = 8) (Harrer et al., 2021). We conducted a Thompson’s regression test as the preferred method to test for funnel plot asymmetry when heterogeneity is high (Jin et al., 2014; Thompson & Sharp, 1999). However, using a funnel plot and regression test to test for asymmetry is otherwise not recommended when k < 10 due to low analytical power (Deeks et al., 2023).

## RESULTS

### Meta-analysis

#### (3) Studies that did not use site size as a biodiversity metric

There is a significant, positive relationship between small mammal diversity, measured as small mammal richness or H’, and *B. burgdorferi* infection prevalence indicating a weak amplification effect. An estimated 1.62 times increase in *B. burgdorferi* prevalence occurs with every unit increase in host diversity (z = 2.96, p = 0.0031) (**Fig. 7.3**). Across studies (k = 6, n = 98), the pooled effect size in a fixed effects model is OR = 1.62 (95% CI [1.18, 2.24]).

#### Publication bias

Publication bias may be present due to studies deviating from a symmetrical distribution indicating an absence of publication bias (**Fig. 2**). The pattern follows a right skewed distribution with most small studies observing null or stronger amplification effects. Studies that would observe stronger dilution effects are potentially missing as indicated by the lack of data points on the left side of the funnel distribution, however, this pattern is not significant (β_0_ = −2.257, p = 0.296).

When contour enhanced to visualize study significance, most small studies fall within the unshaded (white) non-significant (p > 0.1) region on the funnel plot. If hypothetical studies were to be imputed in the empty bottom left corner of the funnel to increase symmetry, they would similarly fall within the non-significant region.

## Appendix Figure Captions

**Figure 1.**
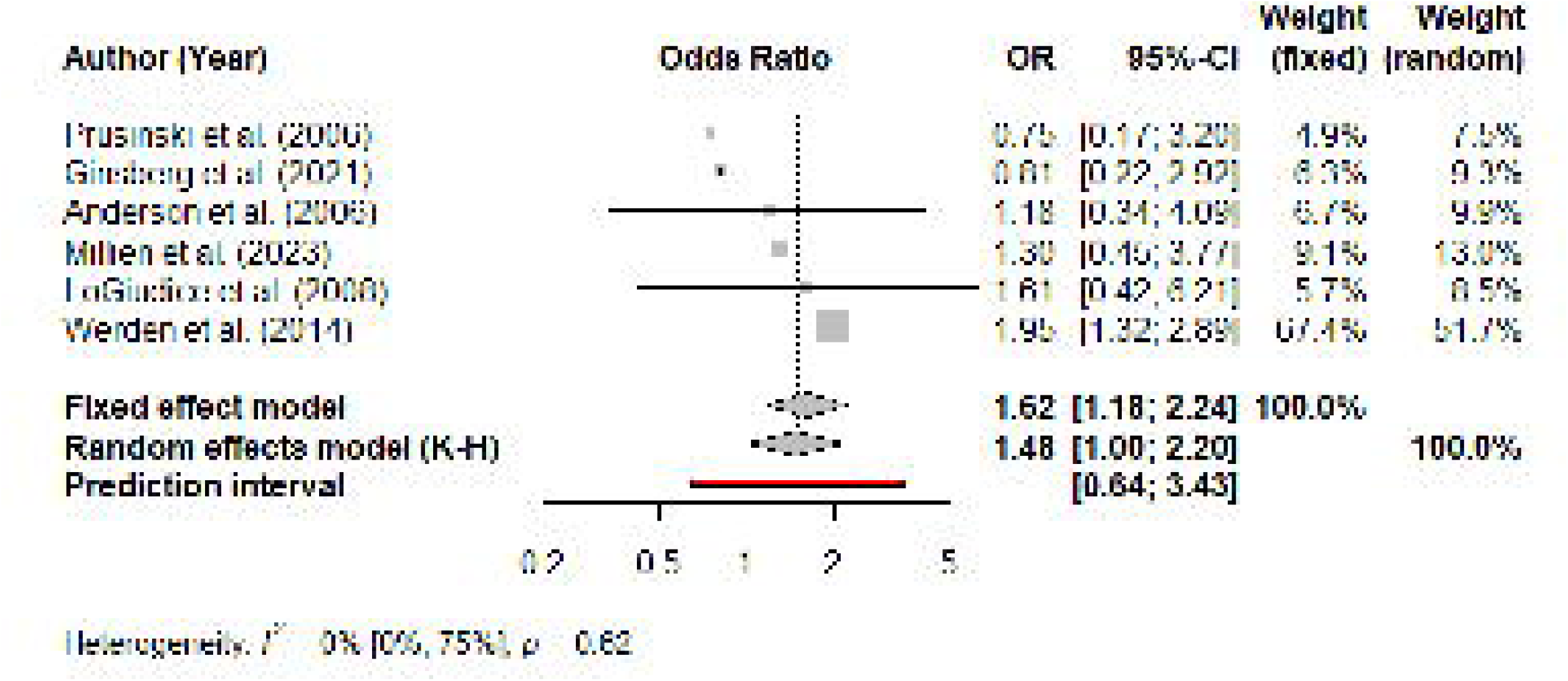
Forest plot showing distributions of odds ratio (with 95% CI bars) for studies describing the relationship between small mammal diversity and disease prevalence using richness or H’ measures. Grey squares are scaled to size according to the weight of that study in the fixed effects model. The solid black vertical line represents the null outcome. The diamonds represent the pooled effect with the corresponding model (fixed effects is the dashed line and random effects is the dotted line). The red horizontal line represents the prediction interval range given the model results.

**Figure 2.**
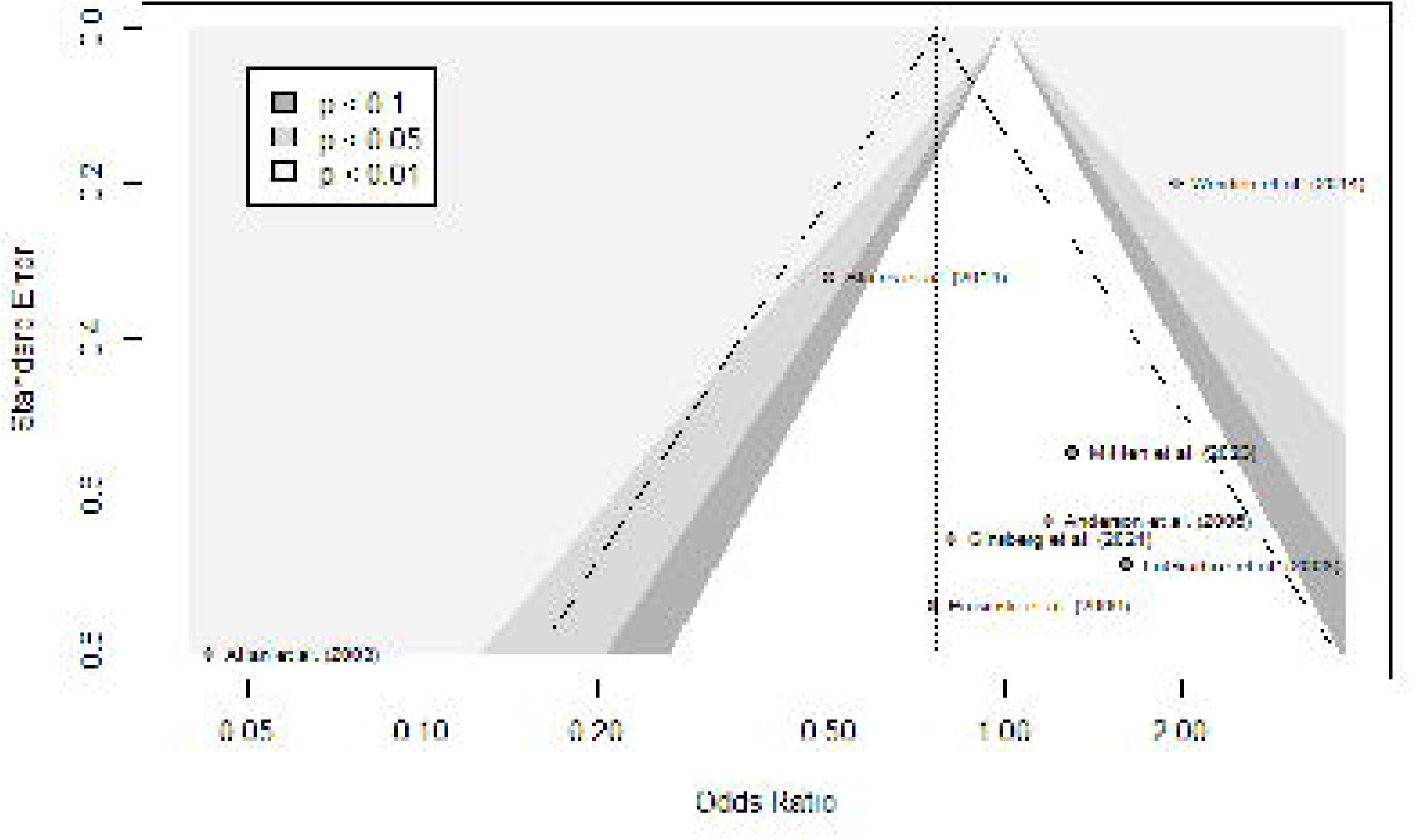
Contoured funnel plot showing the relationship between the observed effect size (OR) and standard error of studies describing the relationship between host biodiversity and *B. burgdorferi* prevalence. Shaded regions (from darkest to lightest) represent varying levels of confidence intervals (p < 0.1, p < 0.05, p < 0.01) around the null (OR = 1). Dotted diagonal lines represent 95% confidence limits around the pooled effect size (dotted vertical line) for each standard error on the y-axis. *Note: three studies (Anderson et al., 2006; Ginsberg et al., 2021; Millien et al., 2023) underwent independent statistical analyses to obtain an effect size and as such these effect sizes were not published in the original paper*.

## Notes

https://github.com/scjpg/lyme-biodiversity-biol4000

### Competing Interest Statement

The authors have declared no competing interest.

